# *Drosophila* co-insulator proteins Pzg and Chro but not CP190 interact with promoter-proximal insulator-binding protein BEAF from a distance

**DOI:** 10.64898/2026.07.26.740821

**Authors:** Lina Shi, Oluwatobi T. Ojemakinde, Craig M. Hart

## Abstract

Chromosomes in *Drosophila melanogaster* are organized into distinct topologically associated domains delimited by boundaries bound by insulator proteins. The insulator protein BEAF (Boundary Element-Associated Factor of 32kDa) plays roles in both chromatin organization and transcriptional regulation, yet its precise molecular mechanisms remain elusive. To examine the role of BEAF in insulator function we used yeast 2-hybrid assays, pull-down assays with bacterially expressed proteins, and bimolecular fluorescence complementation assays in S2 cells to characterize interactions between BEAF and three co-insulator proteins: CP190, Pzg, and Chro. Our studies pinpoint minimal regions of CP190, Pzg, and Chro that directly interact with BEAF, as well as parts of BEAF that are crucial for its interactions with these co-insulator proteins. Functional analyses in transfected S2 cells revealed distinct regulatory roles for BEAF in association with CP190, Pzg, and Chro. CP190 showed a weak interaction with BEAF, and CP190 bound 2.3 kb upstream of promoter-proximal BEAF could not loop out the intervening DNA to effectively communicate with BEAF for luciferase reporter gene activation. In contrast, more robust interactions were observed between BEAF and both Pzg and Chro. Both could also effectively interact with BEAF from a distance to activate the reporter gene. It is likely that the role of CP190 in long-range insulator interactions is mediated by insulator proteins other than BEAF, although BEAF could help stabilize interactions. On the other hand, we propose that BEAF can directly collaborate with Pzg and Chro to mediate long-range chromatin interactions.

## Introduction

The need to explain how enhancers selectively interact with their target promoters led to the idea of insulator elements that prevent promiscuous enhancer-promoter interactions. Among the first insulator elements to be characterized were the scs and scs’ special chromatin structures from either end of the *Drosophila melanogaster* 87A *Hsp70* locus [1–3], the gypsy insulator located near one end of the *Drosophila* gypsy retrotransposon [4], the HS4 DNase I hypersensitive site from the chicken β-globin locus [5], and the mouse *Igf2*-*H19* imprinting control region insulator [6, 7]. The subsequent development of chromosome conformation capture methods coupled with high-throughput sequencing (Hi-C) found that chromosomes are divided into topologically associated domains (TADs), which are chromatin regions with high internal interaction frequencies that are separated by boundaries that restrict interactions between neighboring domains [8, 9]. High-resolution microscopy techniques have independently confirmed this organization [10–16]. TAD boundaries function as insulators by largely limiting regulatory interactions to their resident TADs while also preventing the spread of chromatin states across boundaries.

Just as enhancers obtain their activity from transcription factors and recruited co-activators, insulators derive their activity from sequence-specific DNA binding proteins and recruited co-insulator proteins. In mammals, the main insulator-binding protein is CTCF [17] which, in conjunction with cohesin, plays important roles in establishing and maintaining TAD boundaries. This often involves loop formation between convergently oriented CTCF sites at opposite boundaries of a TAD [18]. Mutations affecting these boundaries, such as small chromosomal duplications, deletions, or inversions, are associated with genetic disorders [19–22]. In contrast, *Drosophila* TAD boundaries are associated with a variety of insulator-binding proteins. For example, the scs’ insulator is bound by BEAF (Boundary Element-Associated Factor of 32 kDa) [23], the scs insulator is bound by Zw5 (zeste-white 5 or deformed wings, dwg) [24], and the gypsy insulator is bound by Su(Hw) (suppressor of Hairy wing) [25]. Although *Drosophila* has a homolog of CTCF (dCTCF), it is present at a minority of TAD boundaries [26–28]. Other *Drosophila* insulator-binding proteins include GAGA factor, Ibf1, Ibf2, Pita, ZIPIC, Mzfp1, Opbp, and TFIIIC [29–34]. Most of these are insect-specific, with several unique to *Drosophila* species [35–37]. Of the co-insulators recruited by these proteins, the best characterized are CP190 (Centrosomal Protein 190), mod(mdg4)67.2 (one of over 30 isoforms of modifier of mdg4), Pzg (Putzig, also known as Z4), and Chro (Chromator, also known as Chriz) [9, 38–42].

Although insulators and associated proteins have been identified, molecular mechanisms concerning insulator formation and function remain unclear. A common model proposes that insulators interact to organize chromatin into looped domains, facilitating regulatory interactions within individual loops while limiting communication across boundaries [43–45]. In mammals, this model is supported by evidence that cohesin-mediated loop extrusion is blocked by head-to-head oriented CTCF sites in boundaries that bracket a TAD, thereby interacting to anchor looped TADs [46–48]. However, only around 15% of CTCF binding occurs within TAD boundaries, indicating that CTCF alone is insufficient for boundary formation [8]. Furthermore, around 60% of mammalian TAD boundaries coincide with active genes, mostly housekeeping genes. This suggests that transcription can contribute to, but is not essential for, boundary formation [8].

In contrast, evidence indicates boundaries on either side of *Drosophila* TADs do not interact to form loop domains [38, 49]. On the other hand, interactions between insulators mediated by the co-insulator proteins CP190 and Chro have been reported [50–53]. Both co-insulators localize mainly to active chromatin [54–56], and over 75% of *Drosophila* TAD boundaries occur at actively transcribed, predominantly housekeeping, genes [28, 57]. It is possible *Drosophila* TAD boundaries assemble in a somewhat stochastic manner to form active chromatin clusters rather than forming discrete loops by specific pairwise interactions [58, 59]. 3D clustering of housekeeping genes has been observed in mammalian cells [60, 61]. Thus, clustering of active chromatin might play the principal role in organizing *Drosophila* chromatin. Looping mediated by mammalian CTCF may have evolved to provide an extra level of organization absent in flies, possibly because mammalian genomes are larger with a lower gene density [62]. Various combinations of insulator and co-insulator proteins are found at *Drosophila* TAD boundaries [34], with BEAF being the most common insulator protein and CP190 and Chro being the most common co-insulators [9, 28, 49]. Therefore, BEAF is well positioned to provide insight into mechanisms underlying *Drosophila* insulator function.

The *BEAF* gene encodes two 32 kDa isoforms, BEAF-32A and BEAF-32B, that we collectively refer to as BEAF [63]. They only differ in their 80 amino acid N-termini (81 amino acids for BEAF-32A) which both contain one BED zinc finger DNA binding domain [64]. Although both are ubiquitously expressed, BEAF-32B has the dominant DNA binding activity and only BEAF-32B is essential [65, 66]. BEAF subunits interact by a C-terminal BESS domain (BEAF-Su(var)3-7-Stonewall domain [67]), with contributions from an adjacent leucine zipper [68]. Evidence indicates BEAF can form dimers and trimers, and formation of larger complexes cannot be excluded [63]. BEAF is required for the function of a class of insulators [65, 69–73]. Yet, genome-wide mapping found that it usually binds near transcription start sites of housekeeping genes [66, 74, 75]. We previously demonstrated that BEAF preferentially activates housekeeping gene core promoters, possibly by helping to recruit chromatin remodeling complexes and the histone chaperone FACT, and can facilitate enhancer-promoter communication through interactions with specific transcription factors [76–78] [79]. Using the *aurA* promoter from the scs’ insulator, we found that the role of BEAF in insulator function and promoter activation can be genetically separated [80]. This suggests these distinct activities rely at least in part on different interacting partners. In particular, interactions with co-insulators rather than transcription factors might underly the role of BEAF in insulator function.

To better understand the role of BEAF in insulator function, here we further investigate reported interactions between BEAF and the co-insulator proteins CP190, Pzg, and Chro [50, 51, 81]. It should be pointed out that this is an interconnected network: Chro has been shown to directly interact with both Pzg and CP190 [50, 82, 83], and CP190, Pzg, and Chro colocalize on chromatin and co-immunoprecipitate from *Drosophila* protein extracts [30, 38, 58, 84, 85]. Here we map domains responsible for interactions between BEAF and the co-insulators, and then examine whether interaction strength correlates with the ability to interact from a distance using a reporter gene assay in transiently transfected *Drosophila* S2 cells. Despite extensive genome-wide colocalization of BEAF and CP190, we find that they only weakly interact and show little communication from a distance. Interactions of BEAF with Pzg or Chro are more robust, and support stronger communication from a distance. Our results are consistent with BEAF primarily playing a role in recruitment of complexes containing Pzg and Chro, with CP190 being incorporated indirectly through its interaction with Chro and probably neighboring DNA-binding proteins. Together with evidence that BEAF is dispensable after early embryogenesis [65] and that BEAF depletion has little effect on CP190 occupancy at BEAF sites [73], our results support a model in which BEAF contributes to the assembly of insulator complexes but functions redundantly with other factors to recruit complexes containing these three co-insulators to chromatin.

## Materials and methods

### Plasmid construction

For yeast 2-hybrid (Y2H) assays, full-length *pzg* was PCR amplified from pBSSKm-Pzg (*Drosophila* Genomics Resource Center LD15904) and fused to the carboxy end of the GAL4 activation domain (AD) in EcoRI cut pOAD [86], the amino end of the AD in KpnI cut pOAD, the carboxy end of the GAL4 DNA-binding domain (BD) in EcoRI cut pOBD2 [86], or the amino end of the BD in NruI cut pBDC [87]. The same strategy was used to make Y2H plasmids for *Cp190* (DGRC LD02352) and *Chro* (DGRC MIP27283). *BEAF-32B* Y2H plasmids have been described [79].

For pull-down experiments, full-length and partial fragments of *Cp190*, *pzg*, and *Chro* were PCR amplified and cloned into a pET-myc expression vector cleaved with KpnI [77]. The plasmid expressing N-terminal FLAG epitope-tagged BEAF-3B was previously described [68]. Individual domains were deleted from this plasmid using outward-directed primers flanking the sequences to be deleted, with a 15 to 20 bp overlap. Gibson assembly was used after PCR and DpnI digestion to destroy the template plasmid to obtain N-terminal FLAG-tagged domain-deleted proteins. To express individual domains and domain pairs, *BEAF-32B* was excised using EcoRI and replaced with PCR-amplified TEV protease cleavage site plus maltose binding protein (MBP) coding sequences with a 5’ EcoRI site and a 3’ XhoI site. This plasmid was cut with BamHI for insertion of PCR amplified parts of BEAF with a 5’ BamHI site and a 3’ NsiI site to code for parts with an N-terminal FLAG tag and C-terminal MBP to aid with stable protein production [88].

Bimolecular fluorescence complementation assays utilized plasmids encoding fusion proteins with the N-terminal half (nV, amino acids 1-173) or the C-terminal half (cV, amino acids 155-239) of the Venus fluorescent protein [89]. Plasmids utilizing the endogenous *BEAF* promoter with nV or cV on the C-terminus of BEAF-32B (BEAF-32B-nV and BEAF-32B-cV) or, as a negative control, cV on the C-terminus of BEAF-32B with the BESS domain deleted (delBESS-cV) were previously described [77]. CP190, Pzg, and Chro were PCR amplified using appropriate primers and inserted into pPac-DREF-cV cut with BamHI and KpnI or pPac-cV-DREF cut with KpnI and XbaI to place cV on their C-termini or N-termini, respectively. The parent plasmids and the transfection control plasmid expressing mCherry-NLS (SV40 nuclear localization signal on the mCherry C-terminus), which all use a 2.5 kb *Act5C* promoter [90], were previously described [79].

Luciferase assays used firefly and transfection control *Renilla* luciferase plasmids that were previously described [79]. Plasmids for protein expression were made by PCR amplification of *Cp190*, *pzg*, and *Chro*, followed by insertion into the EcoRV site of pPac-G4BD and pPac-G4BD-VP16 [79]. The resulting plasmids have an *Act5C* promoter driving expression of fusion proteins with a GAL4 DNA binding domain on their N-termini, without or with a 78 amino acid VP16 activation domain on their C-termini.

Plasmids were made by Gibson assembly (New England Biolabs, Beverly, MA) and identified by colony PCR. Junctions of the assembled plasmids were confirmed by sequencing. Oligonucleotide sequences used are provided in Supplemental Table S1.

### Yeast 2-hybrid assay

Y2H assays were performed as described [68]. Briefly, plasmids were transformed into *Saccharomyces cerevisiae* strain DDY2937 (*MATα trp1–901 leu2–3,112 ura3–52 his3–200 gal4Δ gal80Δ LYS2::GAL1-HIS3 GAL2-ADE2 met2::GAL7-lacZ;* a kind gift from David Donze). Plating on media lacking tryptophan and leucine (2-dropout or 2D) selected for plasmids. After three days at 30°C, colonies were patched onto 2D and 4D (4-dropout: lacking tryptophan, leucine, adenine and histidine) plates to screen for growth due to reporter gene activation, signifying a physical interaction. One colony from a 2D plate was transferred into 2 ml of 2D liquid medium and incubated at 30°C with shaking at 250 rpm. When cells reached exponential growth, cells were diluted to an optical density at 600 nm of 0.1. Four five-fold serial dilutions were made and 5 μl of each dilution was spotted onto 2D and 4D plates. Plates were photographed after incubating for three days at 30°C.

### Pull-down assay

FLAG-tagged and Myc-tagged proteins were expressed in *Escherichia coli* strain Rosetta at 25°C for 24 hours in autoinduction medium ZYM-5052 (1% N-Z-amine, 0.5% yeast extract, 2 mM MgSO4, 25 mM Na2HPO4, 25 mM KH2PO4, 50 mM NH4Cl, 5 mM Na2SO4, 0.5% glycerol, 0.05% glucose, 0.2% lactose, 100 mg/liter ampicillin, and 34 mg/liter chloramphenicol) and protein extracts were prepared using standard methods [91, 92]. For pull-down assays, extracts containing FLAG-tagged BEAF-32B (or an extract lacking FLAG-BEAF-32B as a negative control) were incubated with anti-FLAG M2 magnetic beads (M8823; Sigma-Aldrich, St. Louis, MO), washed 3x with ice cold 10 mM Tris Cl pH 8.0, 150 mM NaCl, aliquoted into tubes, and mixed with extracts containing Myc-tagged proteins. The volume of Myc-tagged protein extract used was based on protein expression levels determined by Western blot. After incubation and washing, pulled down proteins were detected on Western blots using monoclonal anti-Myc (1:1000 dilution; 9E10; Santa Cruz Biotechnology, Dallas, TX) or affinity-purified anti-BEAF (1:10,000 dilution) [23] with ECL chemiluminescence (RPN2232; Cytiva, Wilmington, DE) using established protocols [77].

### Bimolecular fluorescence complementation (BiFC) assay

BiFC assays were done as previously described [77]. Briefly, *Drosophila* S2 cells were cultured at 25°C in Shields and Sang M3 medium (M3; S8398; Sigma-Aldrich, St. Louis, MO) with 10% fetal bovine serum (FBS; 89510-186; GIBCO, Grand Island, NY) and antibiotic/antimycotic (100x is 100 U/ml penicillin, 0.1 mg/ml streptomycin, and 250 ng/ml amphotericin B; 15-240-062; GIBCO). For transfection, cells were seeded into a 24-well plate at a density of 7 x 10E5 cells per well in 1 ml medium and grown for 24 hours. Transfection was performed by mixing 250 ng cV plasmid, 250 ng nV plasmid, and 10 ng mCherry-NLS plasmid with PBS (pH 7.5) in a final volume of 50 μl, followed by addition of 1μl of TransIT-Insect Transfection Reagent (MIR-6100; Mirus Bio, Madison, WI). After incubation for 20 minutes at room temperature, transfection mix was added to each well with gentle mixing. After 2 days further incubation at 25°C, cells were centrifuged at 5000 g for 30 secs and resuspended in 500 μl M3 medium. 10 μl was transferred to a slide with a Secure-Seal Spacer (13 mm diameter, 0.12 mm deep; S24735; Invitrogen, Carlsbad, CA), covered with a coverslip and visualized using a Leica DM6B fluorescence microscope programmed to capture 8 x12 images. The ImageJ ITCN (GitHub - PMB-KU/CountNuclei: Modified Itcn plugin for ImageJ) plug-in was used to count images for Venus-positive and mCherry-positive transfected cells. Images with less than 10 mCherry-positive cells were excluded, and of the remaining images the middle 70 by mCherry count were used for analysis. Total Venus-positive nuclei were divided by mCherry-positive nuclei, and this was normalized to the BEAF-32B-nV, BEAF-32B-cV positive control. Three biological replicates were done, with positive control replicates being divided by the average for the triplicates to calculate the standard deviation of this control. Differences between experimental and control values were analyzed using a two-tailed Welch’s t-test (*p*-value < 0.05 and < 0.005).

### Luciferase assay

Transfections were done following the BiFC assay protocol. Plasmid DNAs used in the assay comprised 5 ng of pPac-*Renilla* Luciferase plasmid (control), 10 ng of pPac-co-insulator protein plasmid, and either 400 ng (for *Rps12* promoter plasmids) or 80 ng (for *yellow* promoter plasmids) of firefly luciferase (test) plasmid. Assays utilized endogenous S2 cell BEAF protein. Cells were lysed 48 hours post-transfection and luciferase activities measured according to the Dual-Luciferase Assay System standard protocol (E1910; Promega, Madison, WI). Luciferase activity was measured using a GloMax 20/20 luminometer (Promega). To control for transfection efficiency, firefly luciferase activity was divided by *Renilla* luciferase activity for each transfection. This value was then normalized to the BEAF binding site promoter plasmid lacking the 4xGAL4 *UAS* for each set of 6 test plasmids and a given co-insulator protein. Three biological replicates were performed. Differences between experimental and control values were analyzed using a two-tailed Welch’s t-test (*p*-value < 0.05 and < 0.005).

### Data availability

Strains and plasmids are available upon request. Primer sequences, BiFC results, and luciferase results are in Supplemental Tables S1-S5. The authors affirm that all data necessary for confirming the conclusions of the article are present within the article, figures, and tables.

## Results

### Full-length Pzg and Chro physically interact with BEAF-32B by Y2H

Previous studies have demonstrated that the chromodomain protein Chro and the zinc-finger protein Pzg co-immunoprecipitate (co-IP) with BEAF, exhibit colocalization on *Drosophila* polytene chromosomes and by genome-wide ChIP peak comparisons, and directly interact with BEAF in yeast 2-hybrid (Y2H) and pull-down assays [58, 81, 83]. Note that the Chro chromodomain lacks consensus amino acids so does not bind methylated lysines [93], and the zinc fingers of Pzg do not confer specific-specific DNA-binding [41, 94]. The same methods have shown that CP190 and BEAF colocalize and interact [50, 51, 74]. We previously identified proteins that co-IPed with FLAG-tagged BEAF-32B from embryo nuclear protein extracts by tandem mass spectrometry (MS) [78]. Both Pzg and Chro were significantly enriched in our co-IP/MS data, but CP190 was not (Table 1). In this study, we sought to confirm the physical interactions of CP190, Pzg, and Chro with BEAF, refine mapping of interaction domains, and determine if these interactions reflect an ability to interact with BEAF from a distance.

**Table 1.** MS data for co-IP of proteins by FLAG-BEAF-32B.

| Protein | FlyBase ID | No. of peptides | Exp/Ctrl | B&H *p*-value |
| --- | --- | --- | --- | --- |
| Pzg | FBgn0259785 | 38 | 3.11 | 0.0390 |
| Chro | FBgn0044324 | 33 | 2.81 | 0.0845 |
| CP190 | FBgn0000283 | 2 | 0.94 | 0.4132 |

We did Y2H assays with BEAF-32B because it has the dominant BEAF DNA binding activity and is essential [65, 66]. Full-length proteins had the GAL4 activation domain (AD) or the GAL4 DNA-binding domain (BD) fused either to their N- or C-terminus, except the BD was only placed on the N-terminus of BEAF-32B because that is the location of its native DNA binding domain (Figure 1A). We found that Chro with an N-terminal GAL4-BD interacted with BEAF-32B with either an N-terminal or C-terminal GAL4-AD (Figure 1A). However, other combinations did not show interactions between Chro and BEAF-32B (Figure S1). Similarly, we found that Pzg with a C-terminal GAL4-BD interacted with BEAF-32B with either an N-terminal or C-terminal GAL4-AD (Figure 1A). Again, other combinations did not detect interactions between Pzg and BEAF-32B (Figure S1). Interestingly, despite previous reports of interaction between BEAF and CP190, we did not observe any interaction in our Y2H assays (Figure 1A and Figure S1).

**Figure 1.**
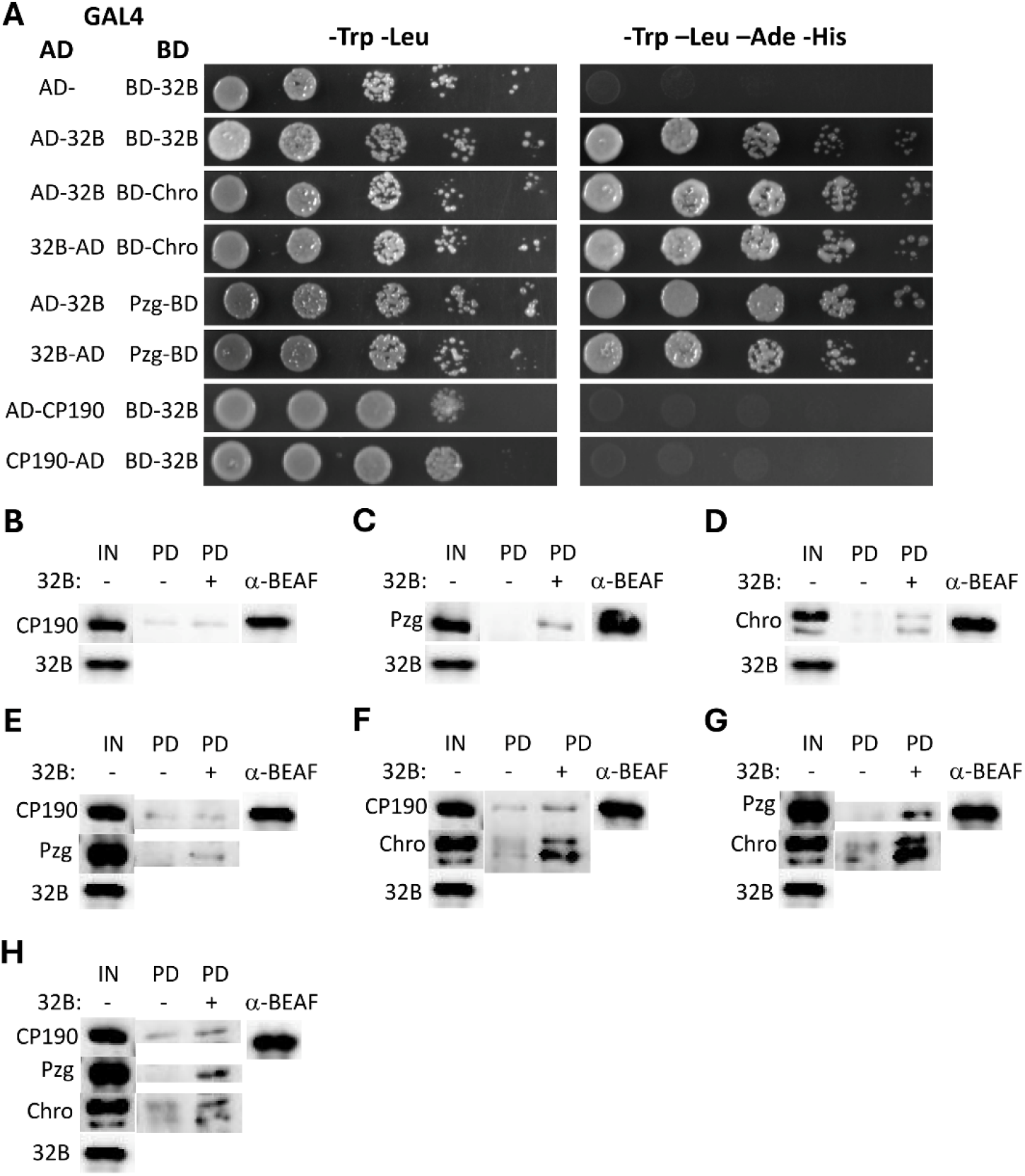
Tests for interaction of BEAF-32B with CP190, Pzg, and Chro. (A) Y2H assays. BEAF-32B had the GAL4 DNA binding domain (BD) fused to its N-terminus or the GAL4 activation domain (AD) fused to either end as indicated by the label position. Chro, Pzg and CP190 had the GAL4-BD or GAL4-AD fused to either end. Combinations showing an interaction are shown, except no interaction between BEAF-32B and CP190 was detected so representative negative results are shown. All other negative results are shown in Supplemental Figure S1. BD-BEAF-32B interacting with empty vector AD (AD-) is shown as a negative control, and interacting with itself (AD-32B) is shown as a positive control. Serial 5-fold dilutions of OD_600_ 0.1 yeast cultures were spotted onto plates. Left panels (-Trp -Leu) show growth on plates selecting for presence of plasmids. Right panels (-Trp -Leu -Ade -His) show growth on plates if reporter genes are activated by BEAF-32B interacting with candidate proteins. (B-H) Pull-down assays with bacterially expressed proteins. Anti-FLAG M2 magnetic beads loaded with FLAG-BEAF-32B were incubated with Myc-tagged proteins of interest prior to pull-down and analysis by Western blot after SDS-PAGE. α-Myc antibody was used to detect Myc-tagged proteins; α-BEAF antibody was used to detect FLAG-BEAF-32B. (B) FLAG-BEAF-32B did not pull down Myc-CP190. (C) FLAG-BEAF-32B pulled down Myc-Pzg, indicating direct interaction. (D) FLAG-BEAF-32B pulled down Myc-Chro, indicating direct interaction. (E) FLAG-BEAF-32B pulled down Myc-Pzg, but not Myc-CP190, when all three were incubated together. CP190 does not affect the interaction between Pzg and BEAF-32B. (F) FLAG-BEAF-32B pulled down Myc-Chro and Myc-CP190 when all three were incubated together. Chro either facilitates the interaction of CP190 with BEAF-32B or acts as a bridge between them. (G) FLAG-BEAF-32B pulled down Myc-Pzg and Myc-Chro when all three were incubated together. Both proteins were more effectively retained by BEAF-32B than in the individual pairwise pull-downs, indicating cooperative interactions between the three proteins. (H) FLAG-BEAF-32B pulled down Myc-Pzg, Myc-Chro, and Myc-CP190 when all four were incubated together, indicating the formation of a complex involving all four proteins. IN, input proteins; PD -, proteins pulled down without FLAG-BEAF-32B; PD +, proteins pulled down with FLAG-BEAF-32B; α-BEAF, FLAG-BEAF-32B from the PD + lane.

### Pzg and Chro, but not CP190, interact with BEAF-32B by pull-down assay

To further explore direct interactions of CP190, Pzg, and Chro with BEAF-32B, we employed pull-down assays using bacterially expressed proteins. N-terminal FLAG-tagged BEAF-32B was incubated with N-terminal Myc-tagged proteins, using an anti-FLAG antibody for the pull-down. Consistent with our co-IP/MS and Y2H findings, Western blot analysis of the pull-down proteins revealed that CP190 did not interact with BEAF-32B (Figure 1B), whereas Pzg and Chro did (Figure 1C and 1D).

We next investigated whether mixing proteins would facilitate interactions with BEAF-32B. We made pairwise mixes of Myc-tagged protein extracts before incubating them with anti-FLAG magnetic beads prebound with FLAG-tagged BEAF-32B. The result with mixed Pzg and CP190 was similar to the results for either protein alone, showing no interaction for CP190 and interaction of Pzg with BEAF-32B (Figure 1E). However, mixed Chro and CP190 resulted in a weak Western blot signal indicating pull-down of CP190 and a stronger Western blot signal for Chro indicating a better pull-down by BEAF-32B than for Chro and BEAF-32B alone (Figure 1F). This agrees with a previous report that Chro and CP190 interact with each other and form a complex that includes BEAF-32B [50]. Mixed Pzg and Chro gave stronger Western blot signals after pull-down by BEAF-32B than for either protein alone (Figure 1G), showing the formation of a mixed complex. This is in line with previous studies that showed Pzg interacts with Chro [82, 83]. When CP190, Pzg, and Chro were mixed, all three were pulled down by BEAF-32B (Figure 1H). Overall, our results demonstrate that CP190 alone does not interact with BEAF-32B. The interaction of CP190 with BEAF-32B was only observed in the presence of Chro or both Chro and Pzg. Additionally, Pzg and Chro interact with BEAF-32B better together than individually.

### Mapping interaction regions of CP190, Pzg, and Chro with BEAF-32B by pull-down assay

To try to further refine the domains required for interaction with BEAF-32B, parts of CP190, Pzg, and Chro were bacterially expressed as Myc-tagged fusions to test for interactions with FLAG-BEAF-32B by pull-down. Although full-length CP190 did not interact with BEAF-32B in our assays, parts did. CP190 amino acids 2-166, which includes the BTB domain, was identified as the smallest region capable of interacting with BEAF-32B (Figure 2A). C-terminal CP190 sequences did not interact with BEAF-32B in our hands, in contrast to an earlier report that found the CP190 acidic C-terminal domain (amino acids 599-1096) interacted [50].

**Figure 2.**
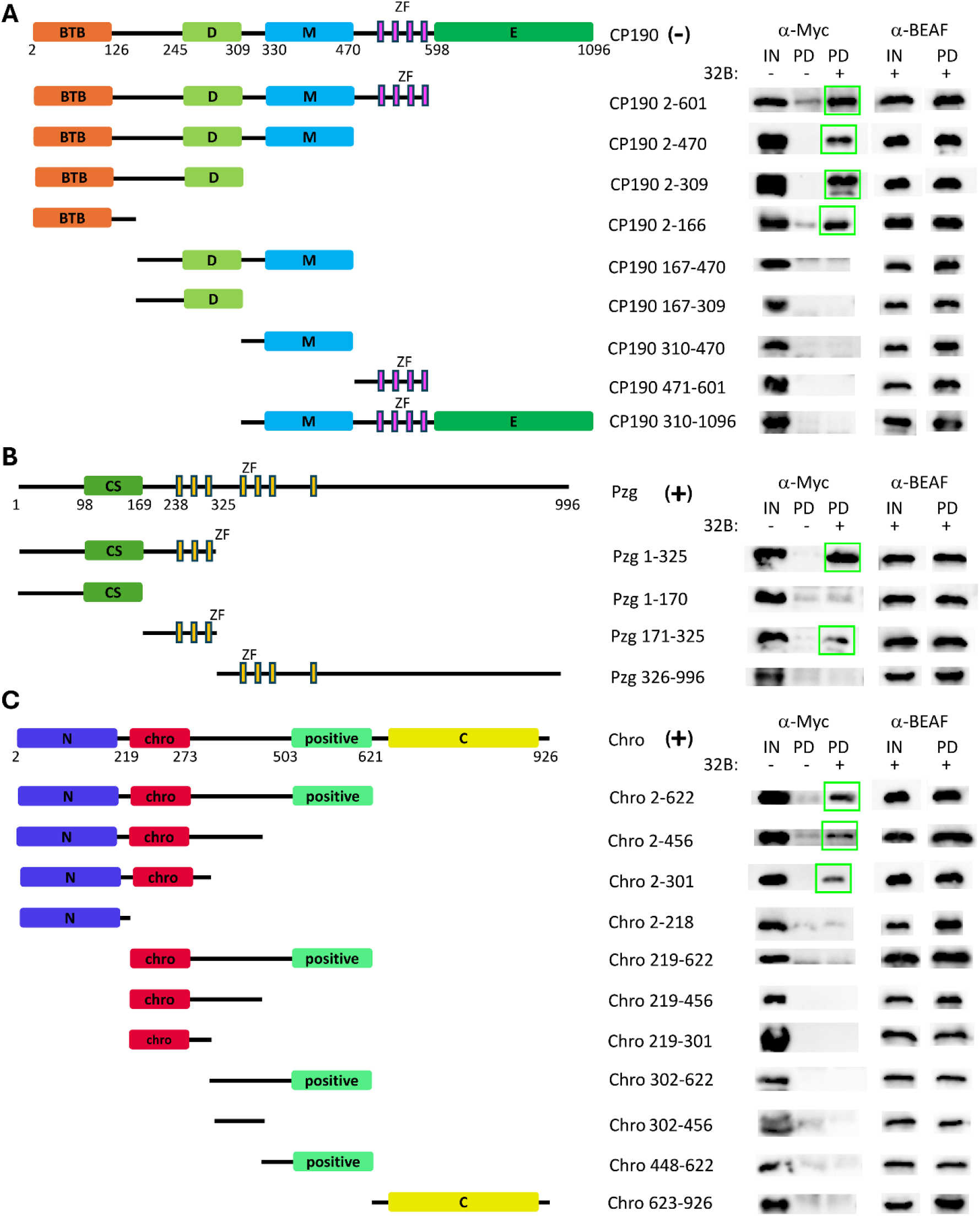
Identification of CP190, Pzg, and Chro domains that interact with BEAF-32B by pull-down assay. Pull-downs were done as described in the legend to Figure 1. (A) Schematic of Myc-tagged CP190 constructs used. CP190 has an N-terminal BTB domain (BTB), an aspartic acid-rich region (D), a centrosomal-targeting domain (M), four C2H2 zinc finger motifs (ZF), and a C-terminal glutamic acid-rich domain (E) [113]. The BTB domain mediated the interaction with BEAF-32B. (B) Schematic of Myc-tagged Pzg constructs used. Pzg is characterized by a conserved sequence (CS) homodimerization domain and seven zinc finger motifs (ZF) [81]. The first three ZF interact with BEAF-32B, but the interaction is stronger if the N-terminus with the CS is also present. (C) Schematic of Myc-tagged Chro constructs used. Chro has an N-terminal conserved domain (N), a Chromodomain (chro), a basic middle sequence (positive), and a C-terminal conserved region (C) [81]. The N-terminal conserved domain plus the chromodomain mediated the interaction with BEAF-32B. Numbers show the approximate limits of the domains, and the first and last amino acids in the indicated truncated proteins. α-Myc lanes detect the Myc-tagged proteins from the input and pull-down without FLAG-BEAF-32B (IN -, PD -) or the pull-down with FLAG-BEAF-32B (PD +). α-BEAF lanes use an α-BEAF antibody to detect FLAG-BEAF-32B from the input and pull-down with BEAF (IN +, PD +). (-) and (+) indicate the interaction results with full-length proteins from Figure 1. Green boxes indicate positive interaction results.

Pzg amino acids 171-325, which encompasses the first 3 zinc fingers, was identified as the minimal region responsible for interaction with BEAF-32B although Pzg 1-325 had an enhanced interaction (Figure 2B). Pzg 1-170 alone did not interact. This is in general agreement with a previous report that found Pzg 1-135 interacted with BEAF-32A, while deleting either the conserved sequences from amino acid 98 to 169 or the first 3 zinc fingers from full-length Pzg eliminated the interaction [81]. Our results suggest Pzg 98-169 is not required, but plays a role in facilitating the interaction of the zinc fingers with BEAF-32B. This region supports homodimerization [95], so facilitation of the interaction could be an indirect effect. Two sets of zinc fingers in a dimer should stabilize binding to a BEAF-32B complex.

The N-terminal region plus the chromodomain of Chro, amino acids 2-301, was identified as responsible for interaction with BEAF-32B (Figure 2C). Neither region alone interacted. This contrasts with previous reports. One found the Chro C-terminal amino acids 601-926 interacted [50], while the other localized the interaction to amino acids 273-621 [81]. The reason for this discrepancy is unclear, although in the next section we confirm the interaction we detected by mapping the region of BEAF it interacts with.

### Mapping interaction regions of BEAF with CP190, Pzg, and Chro by pull-down assay

To identify domains essential for the interaction of BEAF with CP190 2-309, Pzg 1-325, and Chro 2-301, the BEAF-32B DNA binding domain (DBD), Middle region (MID), putative leucine zipper (LZ), and BESS domain of BEAF-32B were individually deleted (Figure 3A). Deleting either the DBD or MID eliminated interactions between BEAF-32B and CP190 2-309 or Chro 2-301, while only deletion of the MID eliminated interactions with Pzg 1-325. Interactions were not affected by deleting either the LZ or BESS domains.

**Figure 3.**
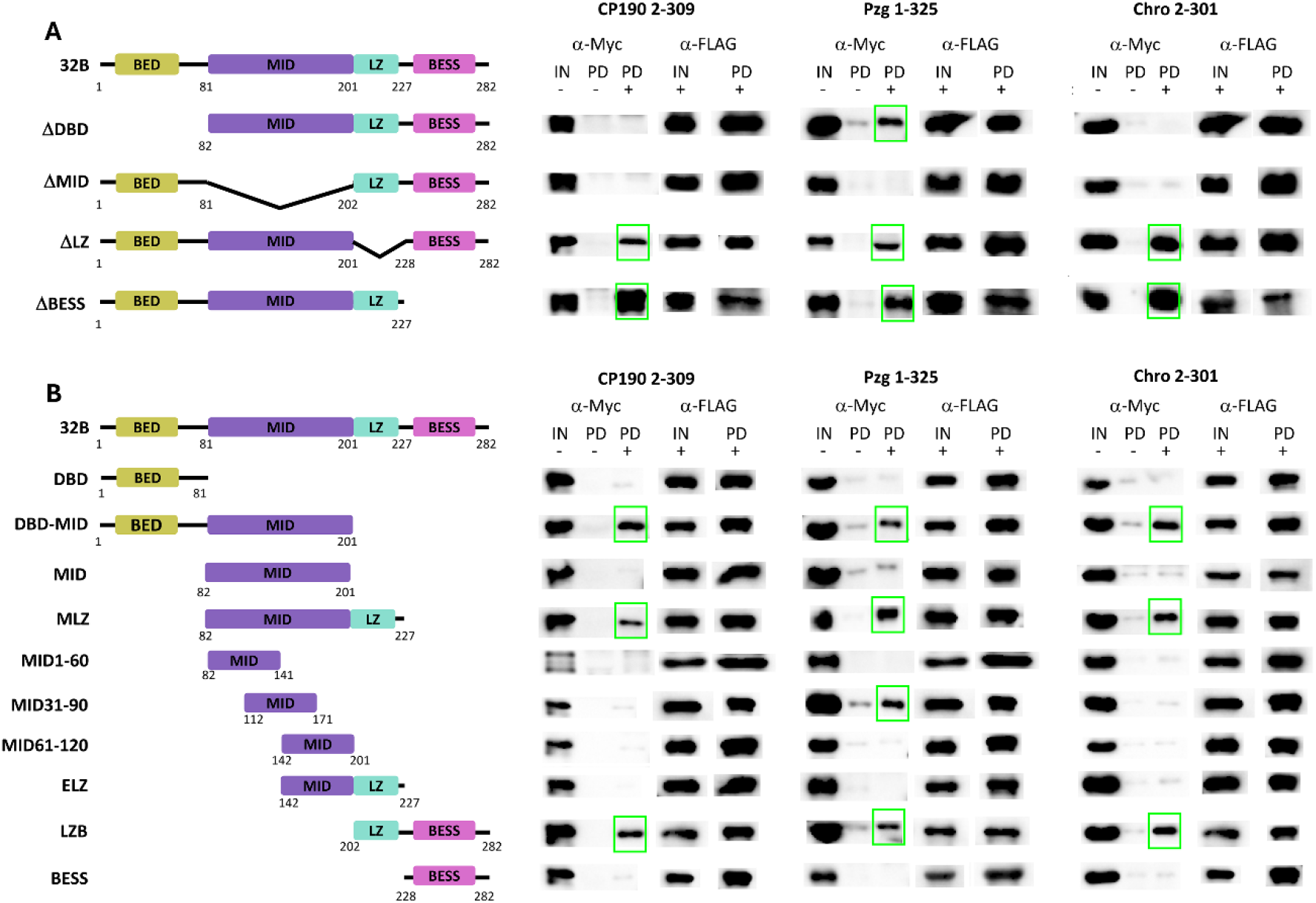
Determination of BEAF-32B domains that interact with CP190, Pzg, and Chro by pull-down assay. Pull-downs were done as described in the legend to Figure 1. (A) Schematic of FLAG-BEAF-32B domain deletion constructs used. Domains in BEAF-32B: BED, N-terminal sequences unique to BEAF-32B, including the DNA-binding BED zinc finger domain; sequences shared with BEAF-32A: MID, intrinsically disordered middle region; LZ, putative leucine zipper domain; BESS, BESS protein interaction domain. Δ signifies the deleted domain. (B) Schematic of BEAF-32B domain constructs used. In addition to an N-terminal FLAG tag, proteins also had a C-terminal maltose binding protein to aid with stable expression in *E. coli*. Note that sequences outside of the N-terminus with the DNA binding BED finger are also present in BEAF-32A. The 120 amino acid MID region is predicted to be intrinsically disordered and was divided into 3 overlapping 60 amino acid parts (MID1-60, first 60 amino acids; MID31-90, amino acids 31-90; MID61-120, amino acids 61-120). Because LZ is only 26 amino acids, to test it without BESS it was extended on its amino side with MID61-120 to give the extended leucine zipper ELZ. Numbers indicate amino acid limits of domains, deleted sequences, and boundaries of truncated proteins. α-Myc lanes detect the indicated Myc-tagged parts of proteins of interest from the input and pull-down without FLAG-tagged BEAF parts (IN -, PD -) or the pull-down with FLAG-tagged BEAF parts (PD +). α-FLAG lanes use an α-FLAG antibody to detect FLAG-tagged BEAF parts from the input and pull-down with BEAF parts (IN +, PD +). Green boxes indicate positive interaction results.

To further refine interaction regions, BEAF was split into individual domains or domain pairs and expressed as FLAG-tagged fusions with the maltose binding protein at their C-termini for stabilization (Figure 3B) [88]. The 120 amino acid MID domain is predicted to be intrinsically disordered (Alphafold Q7JN06) [96]. The last 40 amino acids of MID are poorly conserved among *Drosophila* species, and flies survive if this part of BEAF is deleted [68]. So we also split MID into 3 overlapping parts of 60 amino acids each. Because LZ is only 27 amino acids, it was lengthened with the last 60 amino acids of MID (ELZ).

CP190 2-309, Pzg 1-325, and Chro 2-301 all interacted with MID flanked by DBD or LZ, but did not interact with MID alone. Similarly, all three interacted with LZ when it was flanked by MID or BESS, but not with LZ or BESS alone. This differs from an earlier report that neither Pzg nor Chro interacted with DBD-MID, MLZ, or BESS, but did interact with DBD-MLZ and LZB [81]. We found that Pzg 1-325 showed an additional interaction with MID31-90. Although it was previously reported that BEAF amino acids 140 to 157 (BEAF-32B coordinates) interact with CP190, which correspond to amino acids 59 to 76 of the 120 amino acid intrinsically disordered MID domain [51], CP190 2-309 did not interact with BEAF M31-90, M61-120, or ELZ (which has amino acids 61-120 of MID) in our hands. The simplest interpretation is that MID is critical for interactions with CP190 2-309, Pzg 1-325, and Chro 2-301, with LZ also contributing. The interaction with Pzg 1-325 is further narrowed down to MID31-90. Both MID and LZ require flanking sequences for proper folding or surface accessibility to facilitate interactions, or alternatively, at least in the case of Pzg 1-325, removal of sequences that interfere with interactions. Further studies are necessary to elucidate the specific contacts by which the identified domains in BEAF and interacting partners form complexes, and how this contributes to BEAF function.

### CP190, Pzg, and Chro interact with BEAF-32B in BiFC assays

To further investigate the interactions involving BEAF-32B, we utilized bimolecular fluorescence complementation (BiFC) in *Drosophila* S2 cells (Figure 4). We fused the amino half of the Venus fluorescent protein (nV) to the C-terminus of BEAF-32B (32B-nV) to use with proteins tagged with the carboxy half of Venus (cV). As a positive control we placed cV on the C-terminus of BEAF-32B (32B-cV), and as a negative control we placed cV on the C-terminus of BEAF-32B lacking the BESS domain (delBESS-cV) [68, 77]. C-terminal tags were used with BEAF-32B because the DNA binding domain is at its N-terminus. Because tag placement can affect reconstitution of Venus, we placed cV on either the N-terminus or C-terminus of the co-insulator proteins. Unlike the Y2H or pull-down results, our BiFC assay detected an interaction between full-length CP190 and BEAF-32B. This showed an orientation effect, with cV-CP190 providing evidence for an interaction with BEAF-32B whereas CP190-cV did not. Both Pzg and Chro tagged with cV on either end gave evidence for an interaction with BEAF-32B, although cV-Pzg showed an orientation effect with a lower fraction of fluorescent cells. It is possible proteins present in S2 cells contributed to these interactions. For instance, our pull-down results indicate that Chro can facilitate interactions between CP190 and BEAF-32B. Also, previously reported interactions between Chro and Pzg [83] and our pull-down results suggest they can interact with BEAF-32B as a complex.

**Figure 4.**
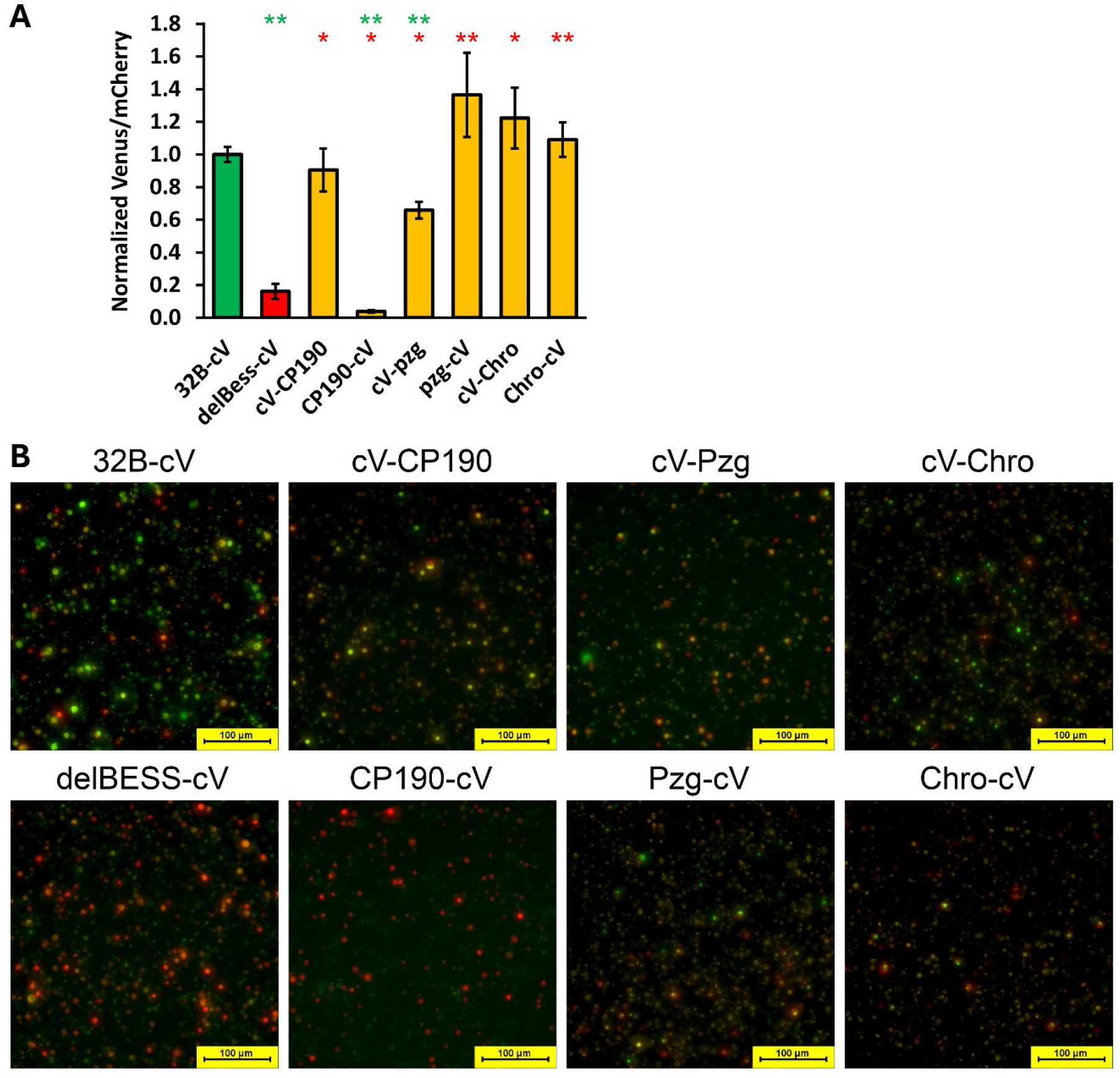
Testing interactions between CP190, Pzg, Chro, and BEAF-32B by BiFC. (A) Bar graph depicting the fraction of cells showing BiFC of cV-tagged proteins with BEAF-32B-nV, normalized to BEAF-32B-cV with BEAF-32B-nV. The cV label position indicates whether proteins were tagged on their N- or C-termini. Data for Venus-positive and mCherry-positive nuclei were derived from 70 images per sample per experiment. Venus-positive nuclei were divided by the mCherry-NLS transfection control prior to normalizing. BEAF-32B-delBESS-cV with BEAF-32B-nV was used as a negative control, and a normalized value less than 0.2 was considered as showing no interaction. Error bars indicate standard deviation (n = 3). Single and double asterisks indicate a *p*-value < 0.05 and < 0.005, respectively, for results compared to 32B-cV (upper, green asterisks) or delBESS-cV (lower, red asterisks; two-tailed Welch’s t-test). All three co-insulator proteins show interactions with BEAF-32B for at least one tagged protein. (B) Representative images for the indicated proteins with 32B-nV. Venus fluorescence is shown in green and mCherry fluorescence is shown in red. Yellow results from overlapping green and red signal. All images were captured using identical settings. Bars represent 100 μm. delBESS, BEAF-32B lacking the BESS domain; cV, carboxy half of Venus; nV, amino half of Venus. Values of the results are given in Supplemental Table S2.

### Pzg and Chro, but not CP190, interact with BEAF from a distance

CP190 and Chro have been implicated in long-range DNA interactions with BEAF [50, 51], and BEAF, Pzg, and Chro frequently co-localize in interbands of *Drosophila* polytene chromosomes [83]. These observations combined with looping models of insulator function prompted us to use a reporter gene assay to investigate whether BEAF interacts from a distance with these co-insulator proteins. Luciferase assays were conducted after transiently transfecting S2 cells as previously described [77] (Figure 5D), modified by replacing the four tandem Sry-δ binding sites with four tandem GAL4 *UAS* sequences [79] (Figure 5A). The assays utilize endogenous BEAF. Test plasmids have a high-affinity BEAF binding site from the scs’ insulator, or as a control a mutated site that BEAF does not bind, immediately upstream of a minimal promoter driving firefly luciferase expression. Plasmids either lack or have four tandem GAL4 *UAS* sequences, either adjacent to the BEAF (or mutated BEAF) binding site (promoter proximal) or 2.3 kb upstream (promoter distal) using phage lambda DNA as the spacer. Transfections also included a plasmid expressing a co-insulator protein with an N-terminal GAL4 DNA binding domain without or with a C-terminal VP16 activation domain (VP16-AD), and a plasmid expressing *Renilla* luciferase under control of the *Act5C* promoter to normalize for transfection efficiency (Figure 5B, C). The firefly to *Renilla* luciferase ratios were further normalized by dividing by the value for the plasmid with a BEAF binding site but no *UAS* sites. The VP16-AD [97, 98] was used because it was not known if the co-insulator proteins alone would provide promoter activation.

**Figure 5.**
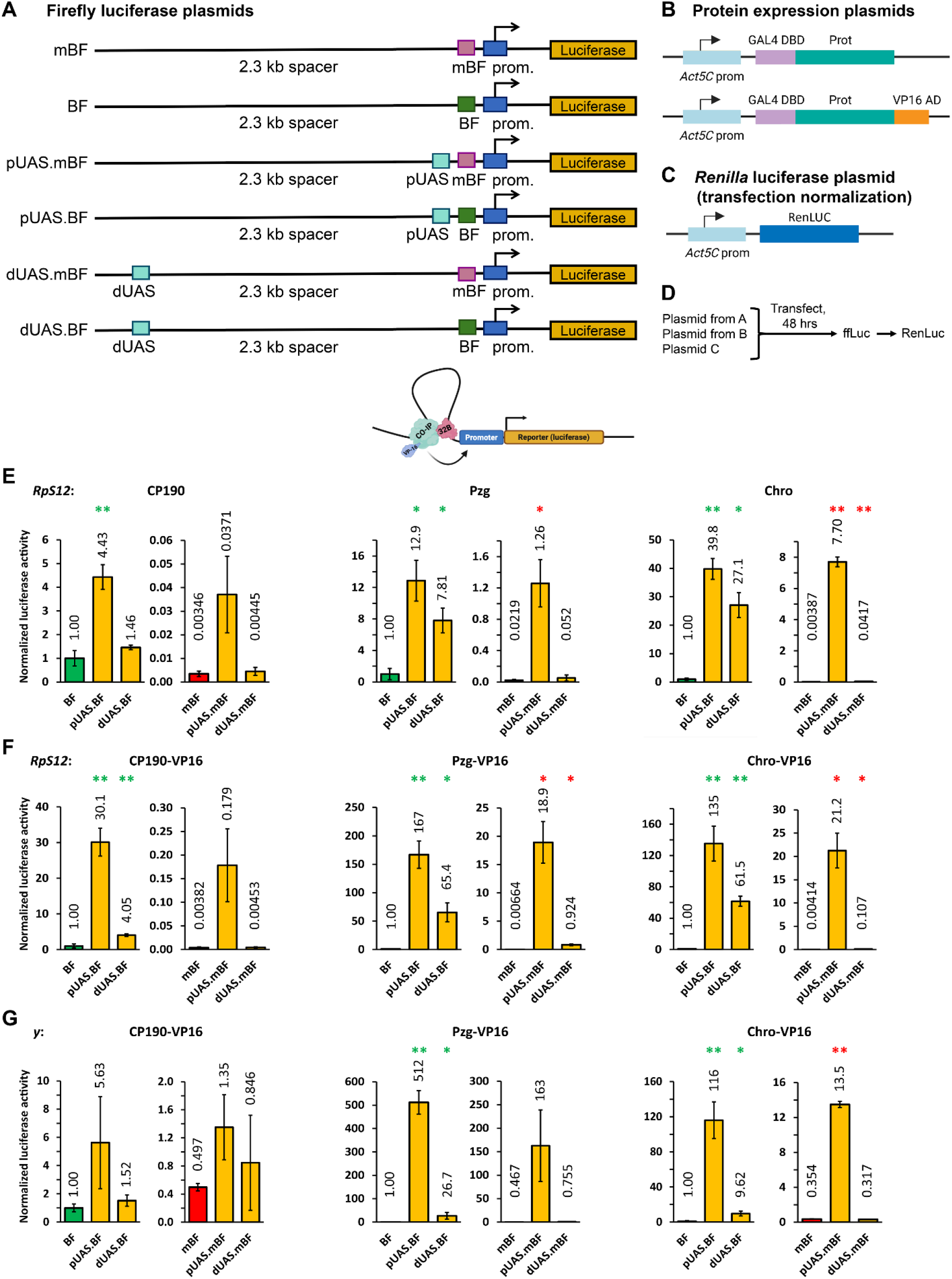
Luciferase reporter gene assay for interaction from a distance between promoter-proximal BEAF and upstream co-insulator proteins. (A) Schematics of the six promoter-firefly luciferase plasmids. Plasmids contain a promoter-proximal BEAF binding site (BF) or mutated BEAF binding site (mBF). When present, 4x GAL4 *UAS* sequences are placed adjacent to (proximal; pUAS) or 2.3 kb (distal; dUAS) upstream of BF or mBF. Plasmids have a 100 bp *RpS12* or 140 bp *y* promoter. mBF plasmids serve as controls for activation without BEAF binding. BF plasmids test for cooperative activation by BEAF and test proteins bound to the pUAS or dUAS sites, with activation from dUAS sites presumably being mediated by looping out the intervening spacer DNA (bottom cartoon; CO-IP, co-insulator protein; 32B, BEAF-32B). (B) Schematics of the protein-expression plasmids, showing the location of the GAL4 DNA binding domain (DBD) and, when present, VP16 activation domain (AD). Expression is driven by a 2.5 kb *Act5C* promoter. (C) Schematic of the *Renilla* luciferase (normalization) plasmid which also uses the *Act5C* promoter. (D) Outline of the experimental protocol. S2 cells were transiently transfected with a firefly luciferase plasmid, a plasmid expressing a co-insulator protein with an N-terminal GAL4 DBD without or with a C-terminal VP16 AD, and the *Renilla* luciferase plasmid. After 48 hours, cells were lysed and assayed first for firefly, and then *Renilla* luciferase activity. (E) Results for the *RpS12* promoter and proteins lacking the VP16-AD. (F) Results for the *RpS12* promoter and proteins with the VP16-AD. (G) Results for the *y* promoter and proteins with the VP16-AD. For each transfection, firefly luciferase activity is divided by the *Renilla* luciferase activity. Then, for each candidate protein, this value is normalized by dividing by the result for the plasmid with the BEAF binding site and lacking the 4x *UAS* sites. Because the normalized values are so low for mBF plasmids with the *RpS12* promoter, BF and mBF plasmid results are plotted separately with different y-axis scales. Error bars show standard deviation for 3 biological replicates. Single and double asterisks indicate a *p*-value < 0.05 and < 0.005, respectively, from two-tailed Welch’s t-tests. Results with BF and UAS sites are compared to BF alone (green asterisks, left plots). Results with mBF and UAS sites are compared to mBF alone (red asterisks, right plots). Values of the results are given in Supplemental Tables S3-S5. See text for details, but results are consistent with all three co-insulators interacting with BEAF from the proximal location, but only Pzg and Chro interacting with BEAF from the distal location. Values of the results are given in Supplemental Tables S3-S5.

Because BEAF usually localizes near housekeeping promoters, we started with a 100 bp *RpS12* housekeeping core promoter. Consistent with previous results [76, 77], BEAF robustly activated this promoter (Figure 5E, F). Without the VP16-AD (Figure 5E), promoter-proximal activation without BEAF binding was 10.7x, 57.5x, and 1990x for CP190, Pzg, and Chro, respectively. Due to variability between triplicates the result had a *p*-value higher than 0.05 for CP190 (two-tailed Welch’s t-test *p* = 0.0687, 0.01917, and 0.0005 for CP190, Pzg, and Chro, respectively). From the promoter distal location, only Chro gave significant activation without BEAF (10.8x, *p* < 0.0001) although this activation was much less than from the promoter-proximal position. Together with BEAF, all three co-insulators activated from the promoter-proximal position (4.43x, 12.9x, and 39.8x with *p* = 0.0015, 0.0136, and 0.0027 for CP190, Pzg, and Chro, respectively). Only Pzg and Chro activated from the distal position with promoter-proximal BEAF (1.46x, 7.81x, and 27.1x with *p* = 0.1237, 0.0124, and 0.0087 for CP190, Pzg, and Chro, respectively). These results indicate that promoter-proximal recruitment of all three co-insulators can activate the *RpS12* promoter, and stronger activation is obtained together with BEAF. However, looping from the distal position to activate with promoter-proximal BEAF only occurred with Pzg and Chro. Although Chro activated from a distance without BEAF, activation was stronger with promoter-proximal BEAF (27.1x versus 10.8x).

To try to get stronger activation, we added the VP16-AD to the co-insulators (Figure 5F). Promoter-proximal binding without BEAF provided stronger activation than without the VP16-AD (46.75x, 2850x, and 5130x with *p* = 0.0595, 0.0123, and 0.0102 for CP190-VP16, Pzg-VP16, and Chro-VP16, respectively). Once again CP190 (now with the VP16-AD) did not activate from the distal position in the absence of BEAF binding, but now both Pzg-VP16 and Chro-VP16 did (1.19x, 124x, and 25.8x with *p* = 0.5787, 0.0100, and 0.0153 for CP190-VP16, Pzg-VP16, and Chro-VP16, respectively). Together with BEAF, all three activated from the promoter-proximal position (30.1x, 167x, and 135x with *p* = 0.0025, 0.0037, and 0.0028 for CP190-VP16, Pzg-VP16, and Chro-VP16, respectively). All three also activated from the distal position with promoter-proximal BEAF, although activation by CP190-VP16 was modest compared to the other two proteins (4.05x, 65.4x, and 61.5x with *p* = 0.0029, 0.0218, and 0.0037 for CP190-VP16, Pzg-VP16, and Chro-VP16, respectively). This is largely consistent with results without the VP16-AD. Distal Pzg-VP16 and Chro-VP16 gave significantly higher overall firefly luciferase activity with promoter-proximal BEAF than without, suggesting communication by looping. Evidence for interaction between BEAF and distal CP190-VP16 is much weaker.

We previously found that promoter-proximal BEAF activates housekeeping core promoters, but not regulated core promoters especially if a TATA box is present [76]. To reduce possible confounding effects of promoter activation by BEAF, we used the VP16-AD fusion proteins with the developmentally regulated, TATA-containing *yellow* (*y*) core promoter (Figure 5G). BEAF did not significantly activate this promoter (2.01x, 2.14x, and 2.83x with *p* = 0.0849, 0.1637, and 0.1926 for CP190-VP16, Pzg-VP16, and Chro-VP16, respectively). Without BEAF binding, Pzg-VP16 and Chro-VP16 activated from the promoter-proximal location although due to variability between replicates for Pzg-VP16 the *p*-value was greater than 0.05 (2.72x, 348x, and 38.1x with *p* = 0.0834, 0.0663, and 0.0002 for CP190-VP16, Pzg-VP16, and Chro-VP16, respectively). None of the three co-insulators with the VP16-AD activated from the promoter-distal location in the absence of BEAF binding (1.70x, 1.61x, and 0.897x with *p* = 0.4657, 0.1843, and 0.1697 for CP190-VP16, Pzg-VP16, and Chro-VP16, respectively). With promoter-proximal BEAF binding, only Pzg-VP16 and Chro-VP16 provided significant activation when bound adjacent to BEAF (5.63x, 512x, and 116x with *p* = 0.1321, 0.0032, and 0.0033 for CP190-VP16, Pzg-VP16, and Chro-VP16, respectively) or 2.3 kb upstream (1.52x, 26.7x, and 9.62x with *p* = 0.1422, 0.0421, and 0.0269 for CP190-VP16, Pzg-VP16, and Chro-VP16, respectively). Taken together, the main conclusion from our luciferase assays is that distally bound Pzg and Chro can loop to interact with promoter-proximal BEAF, while CP190 does this poorly at best.

## Discussion

Understanding insulator function requires identifying proteins involved and their interactions. Various insulator-binding and co-insulator proteins have been identified in *Drosophila*. Here we focused on characterizing interactions between proteins most frequently found in TAD boundaries, the insulator-binding protein BEAF with the three co-insulator proteins CP190, Chro, and Pzg. Although all three co-insulators directly interact with BEAF, our results show that these interactions differ in strength and functional consequence.

Turning first to CP190, our results indicate that both the middle region and the leucine zipper of BEAF contribute to interacting with the CP190 BTB domain. But this interaction is weak in the context of full-length proteins in the absence of additional *Drosophila* proteins such as Chro. The CP190 BTB domain also interacts with the dCTCF, Su(Hw), and Pita insulator proteins [99–101], while other or unknown regions of CP190 interact with the ZIPIC, Opbp, Ibf1, and Ibf2 insulator proteins [29, 30, 32]. Since various combinations of insulator-binding proteins often cluster near transcription start sites in TAD boundaries [28, 34, 57], CP190 recruitment is likely supported by multiple redundant interactions. Support for this comes from finding RNAi-mediated knockdown of BEAF or deleting the CP190 interaction domain from Pita has minor effects on recruitment of CP190 to chromatin [73, 99]. We propose that BEAF plays a supporting role in CP190 association with chromatin, with neighboring insulator proteins being more important for CP190 recruitment. Likewise, while CP190 colocalizes with most BEAF ChIP peaks, CP190 also occupies many additional sites together with proteins such as dCTCF and Su(Hw). This indicates that CP190 participates in multiple, compositionally distinct insulator complexes [38].

In contrast, our results show stronger interactions of BEAF with Pzg and Chro. Pzg and Chro were discovered as interacting proteins that extensively colocalize on polytene chromosomes [41, 55, 83]. We confirmed an earlier report that both also directly interact with BEAF [81]. BEAF interacted more strongly with Pzg and Chro together than individually, paralleling the cooperative effect observed for Chro and CP190. It should be noted that Chro is another protein reported to interact with the BTB domain of CP190 [50]. Like CP190, both the middle region and the leucine zipper of BEAF contribute to interacting with Pzg and Chro. The interaction with Pzg was narrowed down to the central 60 amino acids of the 120 amino acid disordered middle domain, raising the possibility that the leucine zipper relieves inhibition of binding imposed by other parts of the middle domain. The amino termini of both proteins interact with BEAF. For Pzg this includes a conserved domain that mediates homodimerization [95] and the first three of seven zinc fingers. Dimerization might enhance interactions by allowing more zinc fingers to interact with a BEAF complex. For Chro this includes an N-terminal conserved domain and the adjacent chromodomain. As with CP190, redundant interactions are likely involved in Pzg and Chro recruitment to chromatin. For instance, the Opbp insulator protein and M1BP (Motif 1 binding protein) interact with Pzg, Chro, and CP190 [102]. Like BEAF, M1BP is found near many housekeeping gene transcription start sites [103]. Nevertheless, our results suggest BEAF plays a more substantial role in recruiting Pzg and Chro than it does in recruiting CP190.

Like BEAF, these co-insulators could play a role in both insulator function and gene expression. Pzg interacts with a protein complex containing TRF2 (TATA-binding protein related factor 2), the transcription factor DREF (DNA replication factor) and the NURF chromatin remodeling complex [94, 104–106]. The TRF2 complex is important for the function of housekeeping gene promoters, as well as for some regulated genes such as those involved in Notch, ecdysone receptor, and JAK/STAT signaling pathways [94, 105, 107–109]. Chro is implicated in recruiting Pzg to at least some of these promoters [82, 83]. Pzg-Chro complexes also recruit the JIL-1 histone H3 serine 10 kinase to promoters, with JIL-1 spreading into transcribed sequences [83, 110]. In addition, ectopic targeting of Chro or CP190 causes chromatin decondensation [111, 112], and CP190 also interacts with NURF [105]. NURF plays roles in nucleosome organization at both insulators and active promoters [105]. Together, these observations support the idea that BEAF-associated recruitment of Pzg and Chro, and BEAF-assisted stabilization of CP190 binding, contributes not only to insulator function but also to establishing or maintaining an open chromatin environment at active promoters.

A common model proposes that insulators participate in chromatin organization and gene regulation by long distance looping interactions. Many mammalian TADs form loops anchored by pairwise interactions between the TAD boundaries at their ends. Similar TAD loops do not occur in flies, although there is evidence for interactions between insulators. Our test for looping interactions revealed an important functional distinction among the proteins examined. Although CP190, Pzg, and Chro all physically interact with BEAF, only Pzg and Chro effectively communicated with BEAF from 2.3 kb away. The absence of long-range interaction with CP190 agrees with our Y2H and pull-down results and is consistent with the report that CP190 remains associated with BEAF binding sites even after BEAF knockdown [73]. Although this contrasts with previous reports that CP190 can mediate long-range interactions with BEAF [50, 51], our results suggest that direct interaction with BEAF is not sufficient to support communication over this distance.

Results presented here are consistent with results we previously obtained for interactions between BEAF and transcription-associated proteins. BEAF usually binds near housekeeping gene transcription start sites, where it functions in both insulator activity and promoter activation [65, 71, 76, 80]. We found that the transcription factors Sry-δ, ROW, DREF, and Rib all physically interact with BEAF and enhance promoter activation when bound adjacent to BEAF. However, only Sry-δ and ROW, like Pzg and Chro, were able to stimulate transcription when bound 2.3 kb upstream of BEAF [77] [79]. In contrast, DREF and Rib, like CP190, function only locally with BEAF. Thus the ability to interact with BEAF does not necessarily confer the ability to communicate over long distances. Instead, BEAF appears to have at least two mechanistically distinct classes of partners. One class only interacts with BEAF locally. In the case of DREF and Rib, this presumably stabilizes binding for housekeeping gene expression. In the case of CP190, it is likely that BEAF helps stabilize binding mediated by neighboring proteins. If this can involve long-range interactions mediated by these other proteins, that would not be detected in the assay we used because it was sensitized to detect interactions with BEAF. The other class involves interactions that can occur over a distance. In the case of Sry-δ and ROW, this might modulate housekeeping gene expression by enhancer-promoter communication. For Pzg and Chro, this presumably affects insulator function by participating in insulator interactions although direct transcriptional effects cannot be excluded. Understanding how the interactions characterized here contribute to insulator function, and the molecular basis for the ability to make long-range interactions rather than only local, are important goals for future studies.

