## Supplemental Figure S1 for "*Drosophila* co-insulator proteins Pzg and Chro but not CP190 interact with promoter-proximal insulator-binding protein BEAF from a distance"

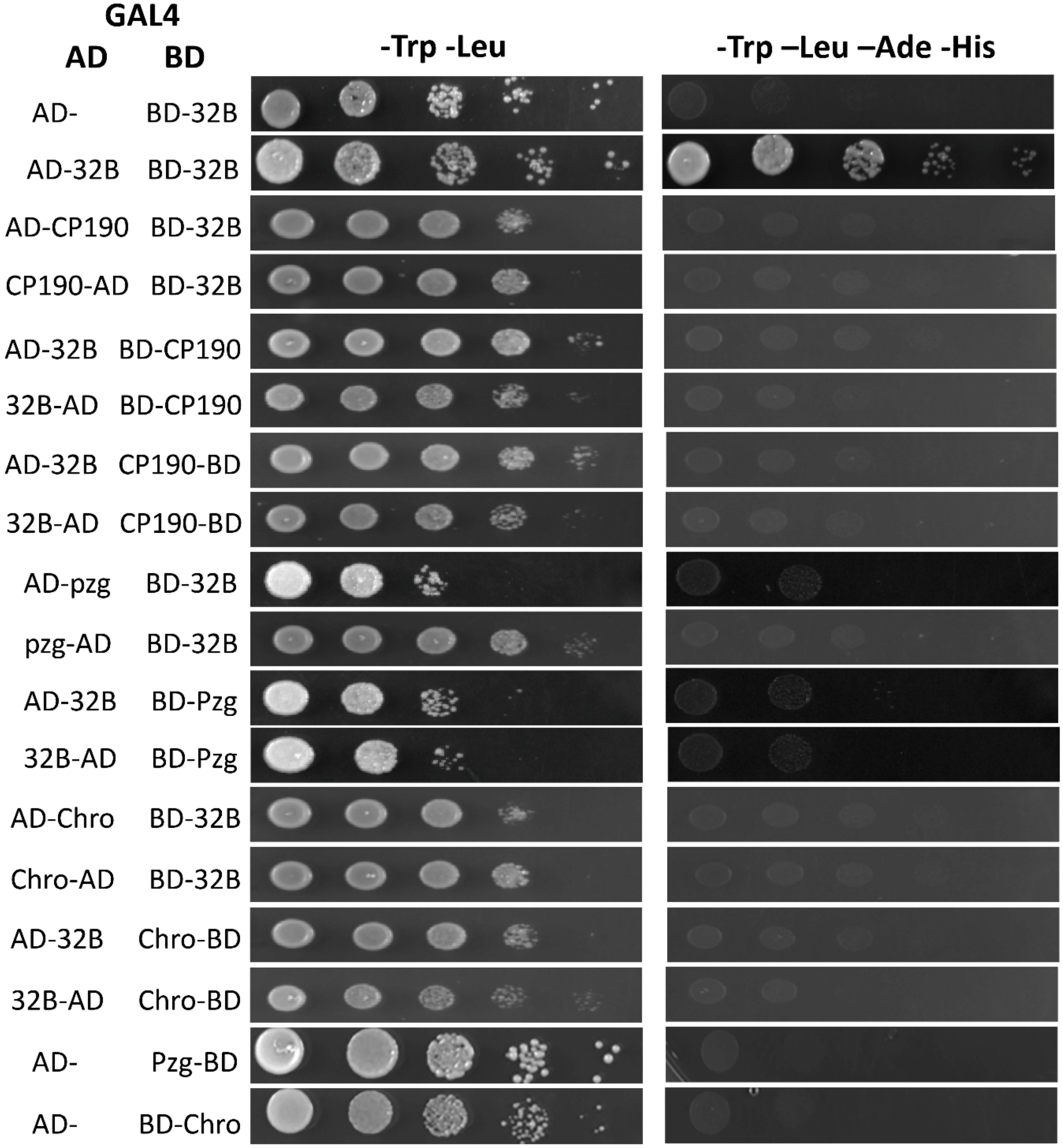


**Figure S1. Tests for interaction of BEAF-32B with CP190, Pzg, and Chro by Y2H: negative results.** BEAF-32B had the GAL4 DNA binding domain (BD) fused to its N-terminus or the GAL4 activation domain (AD) fused to either end as indicated by the label position. Chro, Pzg and CP190 had the GAL4-BD or GAL4-AD fused to either end. Combinations not showing an interaction are shown. BD-BEAF-32B interacting with empty vector AD (AD-) is shown as a negative control, and interacting with itself (AD-32B) is shown as a positive control. Serial 5-fold dilutions of $OD_{600}$ 0.1 yeast cultures were spotted onto plates. Left panels (-Trp -Leu) show growth on plates selecting for presence of plasmids. Right panels (-Trp -Leu -Ade -His) show growth on plates if reporter genes are activated by BEAF-32B interacting with candidate proteins. No BD fusions showed autoactivation with empty vector AD, but only Pzg-BD and BD-Chro are shown because other BD fusions gave negative results with AD fusion proteins, which are shown.
